# Subgenomic satellite particle generation in recombinant AAV vectors results from DNA lesion/breakage and non-homologous end joining

**DOI:** 10.1101/2020.08.01.230755

**Authors:** Junping Zhang, Ping Guo, Xiangping Yu, Derek Pouchnik, Jenni Firrman, Hongying Wei, Nianli Sang, Roland Herzog, Yong Diao, Weidong Xiao

**Author notes:** These authors contributed equally to this work (Junping Zhang, Ping Guo, Xiangping Yu).

## Abstract

Recombinant AAV (rAAV) vectors have been developed for therapeutic treatment of genetic diseases. Nevertheless, current rAAV vectors administered to patients often contain non-vector related DNA contaminants. Here, we present a thorough molecular analysis of the configuration of non-standard AAV genomes generated during rAAV production. In addition to the sub-vector genomic size particles containing incomplete AAV genomes, our results found that rAAV preparations were contaminated with multiple categories of subgenomic particles with either snapback genomes or vector genomes with deletions in the mid regions. Through CRISPR and restriction enzyme-based in vivo and in vitro modeling, we identified that the main mechanism leading to the formation of non-canonical genome particles occurred through nonhomologous end joining of fragmented vector genomes caused by genome lesions or DNA breaks that were generated by the host cell/environment. The results of this study advance our understanding of AAV vectors and provide new clues on improving vector efficiency and safety profile for use in human gene therapy.

## Introduction

Recombinant adeno-associated virus (rAAV) vectors have been widely adopted as a gene delivery tool for basic research as well as a pharmaceutical drug vector for human gene therapy (*1*). The vector genome is constructed by inserting the desired expression cassette and regulatory elements between two flanking copies of inverted terminal repeats (ITR). The ITR functions as the replication origin for AAV vectors and as the packaging signal for the rAAV production process. rAAV vectors are typically produced by transfecting host cells, such as 293 cells, with plasmids coding for the vector while also supplying helper functions by delivering either a helper virus, such as adenovirus, or trans factors, such as rep78 and rep68, rep52/rep40, and VP1, VP2 and VP3. Alternatively, rAAV vectors may also be produced using a non-adenovirus helper or a non-mammalian system with baculovirus.

While rAAV vector preparation can be performed following a standard procedure, it does not result in the production of a homogenous population, even for GMP produced vectors that are used clinically. Previously identified vector related impurities include AAV particles containing plasmid backbone sequence and even host genomic sequences (*2*). Although defective interference particles are known to exist in wild type AAV populations (*3*–*5*), similar particles found in recombinant AAV vector preparations have never been fully characterized due to technical difficulties in obtaining the detailed vector sequences from the entire population. Previously, next generation sequencing (NGS) has been used to profile the rAAV genomic configuration and to perform transcriptomic analysis (*6*). The helicos-based sequencing platform has been used to profile the 3’ end of the rAAV genomes (*7*). However, all of these data have only partial genomic information on the rAAV system. In a separate study, PacBio SMRT sequencing was used to produce more long reads and covered some special categories of rAAV genomes in the vector population (*8*). Here we systemically characterized the molecular state of rAAV vector genomes at a single virus level. In addition, through CRISPR-Cas9 based in vivo modeling, we identified that the host-mediated vector genome lesion/breakage as the main cause for AAV vector subgenomic particles formation.

## Results

### Molecular configuration of subgenome particles in the rAAV population suggested Non-homologous end joining (NHEJ) events during AAV replication and packaging

In order to reveal the molecular state of individual AAV genomes in a population produced by the typical triple plasmid transfection method, we took advantage of the long reads and high accuracy of the PacBio Single Molecule, Real-Time (SMRT) Sequencing platform. From analyzing viral genomes at single virus level from at least 10 different rAAV preparations, including single stranded DNA genome vector as well as self-complementary DNA vectors, the highly heterogeneous rAAV population with only AAV genome DNA sequences was classified within the following categories based on the our analysis of thousands of vector genomes (Fig. S1): 1. Standard rAAV genomes which contained the complete vector sequences including transgene expression cassette and flanking AAV ITRs; 2. Snapback genomes (SBG) which had the left or right moiety of standard duplex rAAV genomes. The SBG was further classified as symmetric SBG (sSBG) or asymmetric SBG (aSBG) according to the DNA complementary state of the top and bottom strands. For sSBG, the top and bottom strands complemented each other. Unlike sSBG, DNA at the bottom strand of aSBG did not match the top strand completely and, therefore, promoted loop formation in the middle region; 3. Incomplement rAAV genomes (ICG), which had an intact 3’ITR and partial AAV genome. These were presumably formed by an aborted packaging process; 4. Genome deletion mutants (GDM), in which the middle region of the AAV genomes were deleted; 5. Secondary derivative genomes (SDG) which were formed by using class 2-4 molecules as the template and the same mechanism to generate the next generation of subgenomic vector molecules of class 2-4.

While typical sSBG configuration may have been the product of a template switch, the existence of aSBG, GDM, and SDG could not be explained by a template switch of the DNA polymerase during AAV replication. Since there were remnant signs of multiple DNA fragments in the aSBG, GDM and SDG, we proposed that NHEJ events had occurred during the AAV replication and packaging processes.

### NHEJ as the mechanism for generating subgenomic particles in an rAAV population

An NHEJ reaction requires the presence of corresponding DNA fragments. The dissection genomic configurations of subgenomic particles of AAV suggested the existence of such fragments. First we tested whether or not NHEJ events could led to the generation of snapback genomes (SBG). We transfected host cells with linear rAAV DNA fragments (Fig. 1A) that were generated though restriction enzyme digestion in the presence of trans elements that complement AAV replication and packaging (Fig. 1). The parent vector plasmid pCB-EGFP-6.4K was oversized for AAV capsids. DNA recovered from vectors prepared using this oversized plasmid primarily consisted of smaller fragments, which were less than 6.4 kilobases in size. In contrast, vectors prepared from smaller vector plasmid pCB-EGFP-3.4K, which falls within the packaging limits for the AAV capsid, mainly produced viral particles with a 3.4kb DNA genome. Interestingly, when linear fragments derived from pCB-EGFP-6.4K ranging from 0.6kb to 3.1kb were used for transfection, the most prominent genomes recovered from the prepared vectors appeared to be resultant of intermolecular nonhomologous end joining (Fig. 1B). Even though intra-molecular DNA joining of the 5’end and 3’end was supposed to be more efficient, the vectors resulting from such reaction were in relatively lower yield. This was most likely because their size was larger than the inter-molecular NHEJ products. The inter-molecular NHEJ products were confirmed to be snapback genomes (SBG, data not shown). More specifically, when vector was prepared using fCB-GFP-2.3K, the vector DNA size from AAV ITR to the breaking points was 1.8kb and 2.3kb respectively. The main vector size shown in the gel were 1.8kb (3.6kNT in single stranded form), 2.3kb (4.6kNT), along with a faint 4.1 kb region (which was annealed from both plus strand and minus strand) which suggested intramolecular joining. Similar observations were obtained for vectors prepared using fCB-GFP-0.6k (0.6kb-1.2kNT, 1.8kb-3.6kNT), fCB-GFP-1.0k (1.0kb-2.0kNT, 1.8kb-3.6kNT), fCB-GFP-1.6k (1.6kb-3.2kNT, 1.8kb-3.6kNT), and fCB-GFP-1.8k (1.8 kb-3.6kNT). The exception was for vectors prepared using fCB-GFP-3.1k, in which we only observed a 1.8kb genome fragment. This is likely because the 3.1kb SBG molecule, which was 6.2kNT, was over the packaging size limit for AAV vectors.

**Fig 1.**
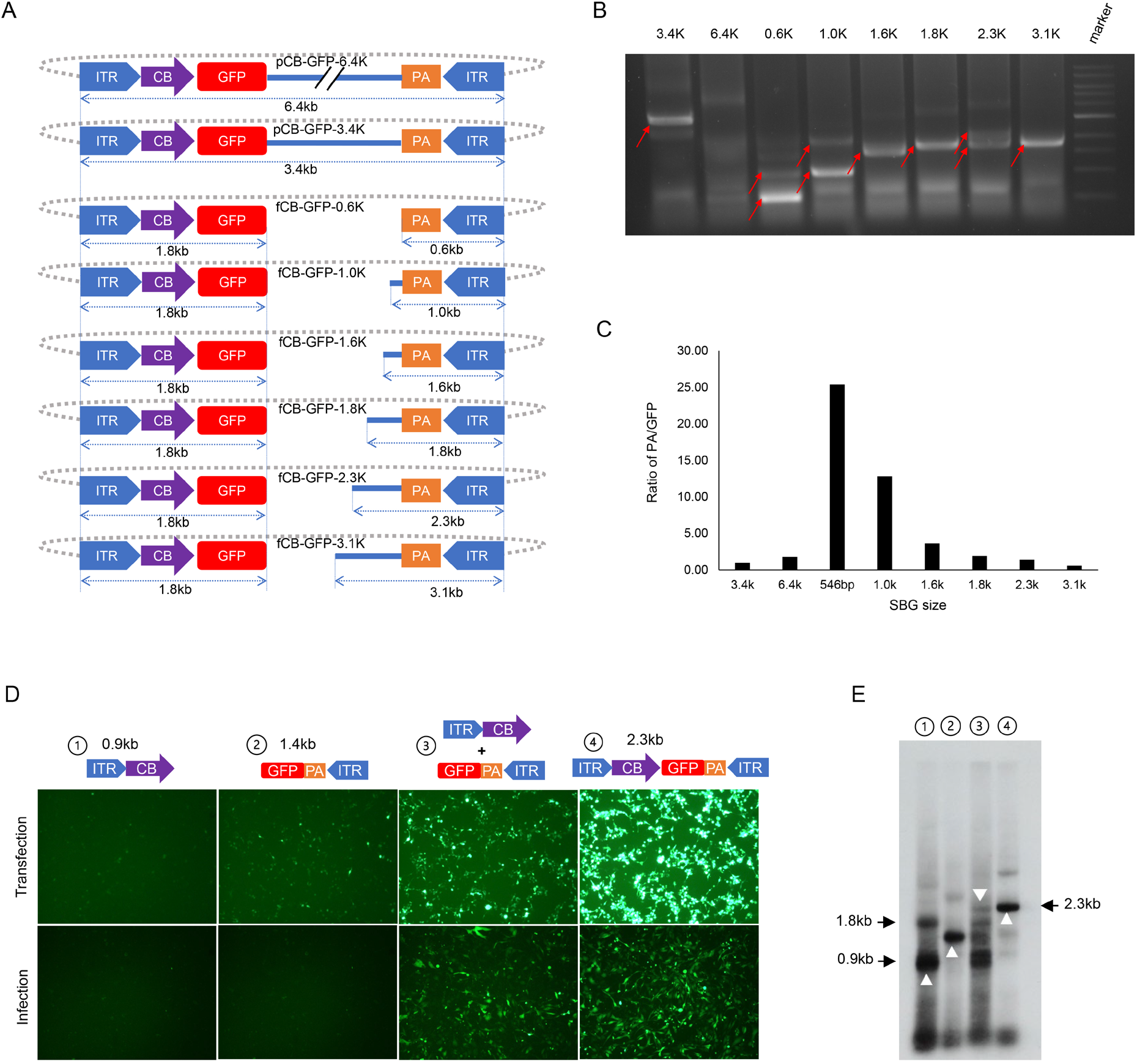
Intermolecular NHEJ is a mechanism leading to the formation of SBG molecules. The parent plasmid 6.4kb pCB-GFP-6.4K was linearized with varying restriction enzymes to obtain linear fragments as shown in A). The plasmid backbone is depicted with a dotted line. Hek293 cells with rAAV packaging helper functions were transfected with DNA fragments in (A), plasmid pCB-GFP-6.4K, or pCB-GFP-3.4K. B) The resulting rAAV vectors in the media were harvested and the DNA in the vectors were extracted and analyzed for genome status using a 1% agarose gel. For simplicity, fragments such as fCB-GFP-0.6K were referred to as 0.6k on top of the gel in B. Red arrows indicate key fragments. The vectors recovered were quantified by qPCR using primers specific for poly A or GFP. The ratio of vectors containing poly A or GFP are shown in C. In pCB-GFP-3.4K, the vector size is 3.4kb. For DNA fragments, the size of 5’ITR-GFP is 1.8kb and the size of the poly A to 3’ITR are indicated as the last three letters in the name. D. E. showed Intermolecular NHEJ is a mechanism leading to AAV genome deletion mutants (GDM). D) Hek293 cells were transfected with ITR fragments containing the CB promoter or GFP gene alone or combined with supplemental helper plasmids for rAAV replication and packaging. The positive control was a 2.3kb intact pCB-GFP-3.4K plasmid. At 3 days post-transfection, the GFP expression was monitored by fluorescence microscopy (mid panel). The harvested vectors were used to transduce GM16095 cells, and the GFP expression was monitored at 24 hours post-infection (bottom panel). E). The vector DNA recovered from panel A was electrophoresed in 1% agarose gel and AAV genomes were detected by Southern blot using an ITR specific probe. Δ indicate key fragments 0.9kb, 1.4kb and 2.3 kb.

When these fragments were used to supply AAV production, we noticed the relative abundance differed among vectors produced. Since the Poly A containing vectors or GFP containing vectors represented different NEHJ reactions, the ratio of these two types of vectors was graphed in Fig. 1C. Based on these results, it was evident that the smaller sized subgenomic particles became more dominant. This suggested that later DNA replication and packaging favor smaller genomes, which may be a major mechanism dictating the abundance of rAAV subgenomic particles. To generate genome deletion mutant (GDM), the fragments representing 5’ ITR and 3’ITR had to be linked together through NHEJ. To demonstrate this, we transfected Hek293 cells with a 5’ ITR fragment carrying the CB promoter and a 3’ITR fragment carrying the GFP gene along with AAV replication and packaging helper plasmids. Interestingly, the combination of these two fragments efficiently regenerated the functional GFP expression. The infectious GFP vectors also regenerated as shown in the transduction assay (Fig. 1D). As presented in Fig. 1E, the anticipated genome size can be observed in the southern blot. This experiment demonstrated that the GDM molecules were produced through the same mechanism that generated the SBG virus (Fig. 1E).

### DNA lesion/nicking is sufficient for generating AAV subgenomic particles

To investigate if a DNA lesion was sufficient to generate SBG molecules, CRISPR-cas9 nickase activity was introduced to the AAV production system (Fig. 2). As presented in Fig. 2B, the in vivo cutting with Cas9 at various positions generated two major SBG molecules corresponding to the cutting sites. Using gRNA9 as an example, Cas9 cutting generated two vectors at 1.5kb and 1.9kb respectively. However, nicking at the top strand by gRNA9 and Cas9-H840A yielded a 1.5kb DNA. Nicking at the bottom strand by gRNA9 and Cas9-D10A yield a 1.9kb DNA. In contrast, cutting of the vector by gRNA4 and Cas9 yielded two main vectors, 0.6kb and 1.2kb in size. The 2.8kb (5.6kNT) SBG vectors that should have appeared was not observed because it exceeded the packaging capacity of AAV particle. Nicking with gRNA4 and Cas9-H840A yielded 0.6kb vector along with its dimer at 1.2kb. On the other hand, nicking at the bottom strand by gRNA4 and Cas9-D10A yielded no major bands since the theoretical 2.8kb(5.6kNT) SBG vectors are oversized for AAV capsids. Similar results were obtained from gRNA13 induced nicking or cutting. The other exception was when NHEJ product replication was overwhelmed by their relatively smaller companion fragments. In this case, the corresponding larger DNA was not greatly diminished, i.e. gRNA5 and gRNA10. This result suggested that DNA lesion/nicking was sufficient to generate DNA fragments that can lead to the creation of subgenomic particles. There was a clear strand selection, in which the nicking site and its 3’ end ITR formed snapback molecules.

**Fig. 2.**
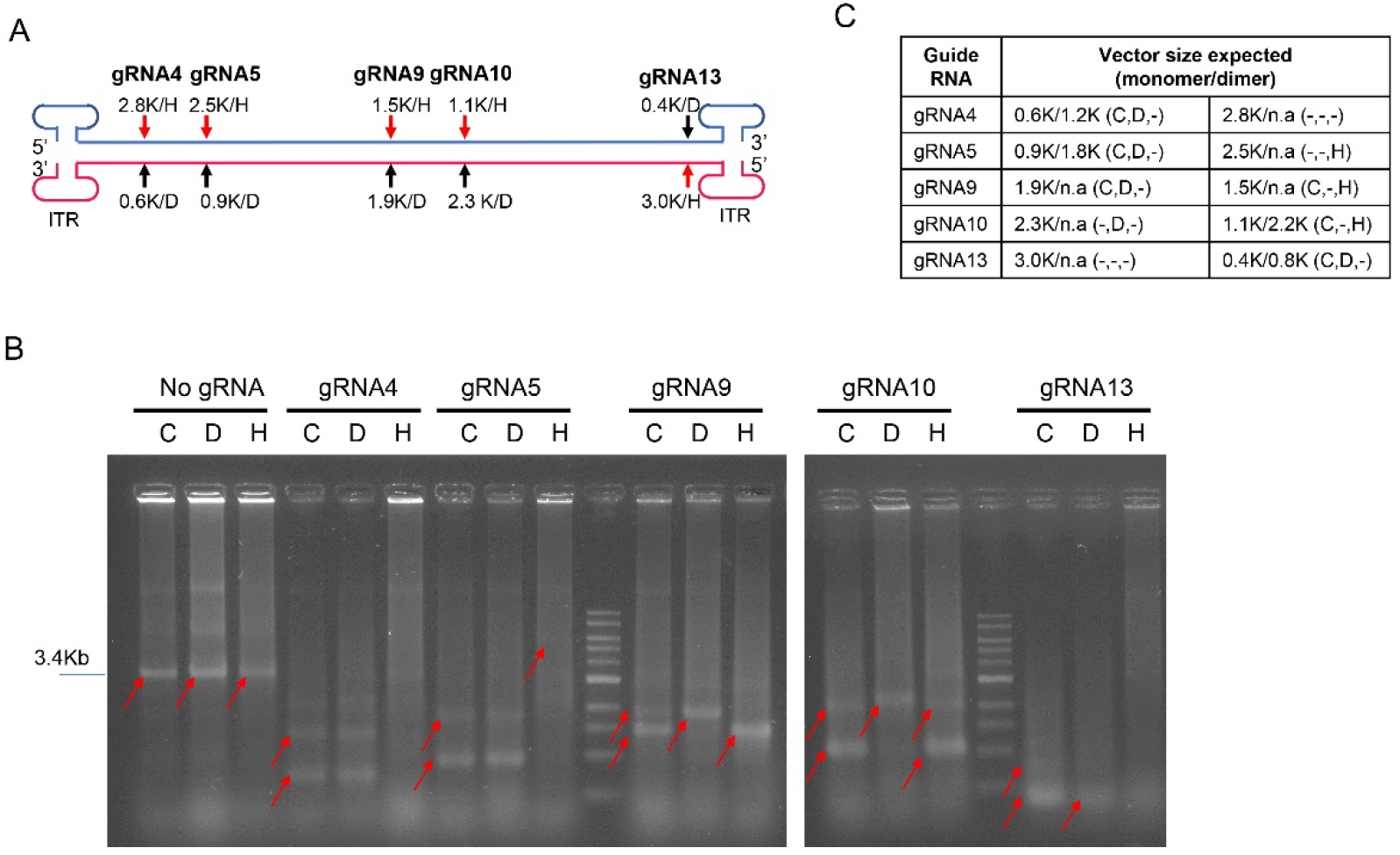
Intra-host cells vector DNA Lesion is sufficient for SBG formation. A) Illustration of gRNA sites in pCB-EGFP-3.4K for Cas9 nicking or digestion. Hek293 cells were transfected with plasmid pCB-EGFP-3.4K for vector production in the presence of cas9 or cas9 mutants (H840A or D10A) and corresponding guide RNA. B) The resulting vector DNA was electrophoresed in native agarose gel with EB staining. C) Cas9-double cut, D: D10A-nicking H: H840A-nicking. H stands for Cas9-H840A nicking. D stands for Cas9-D10A nicking. C stands for Cas9 cutting. The table summarized the potential DNA sizes that can be generated by nicking or cutting. The actual observed bands are summarized in the brackets. - indicates “not observed”.

### Cellular DNA damage in events led to subgenomic molecule formation

We further hypothesize that intracellular DNA damage events may lead to subgenomic DNA formation. As shown in Fig. 3, hydrogen peroxide was added to examine the effects of this DNA damage reagent on AAV production. Corresponding to an increased concentration of hydrogen peroxide, the recovered rAAV vectors appeared as smears that were smaller in size compared to the standard AAV vectors. At 200mM hydrogen peroxide, the majority of vector DNA detected were small subgenomic DNA particles (Fig. 3A). We prepared a library of the recovered DNA from these vectors and performed DNA sequencing using the PacBio SMRT sequencing platform. More than 50,000 genomes were sequenced. The majority of these sequences were not AAV vector related and appeared as short DNA fragments. Those sequences aligned to the initial vector appeared to be heavily fragmentated (Fig. 3B). In addition, SBG produced from the initial vector were recovered in the sequencing as well (Fig. 3B). Some of these molecules were found to contain portions of the plasmid backbone. These results showed that global DNA damage events can ruin recombinant AAV production and lead to the production of subgenomic particles.

**Fig. 3.**
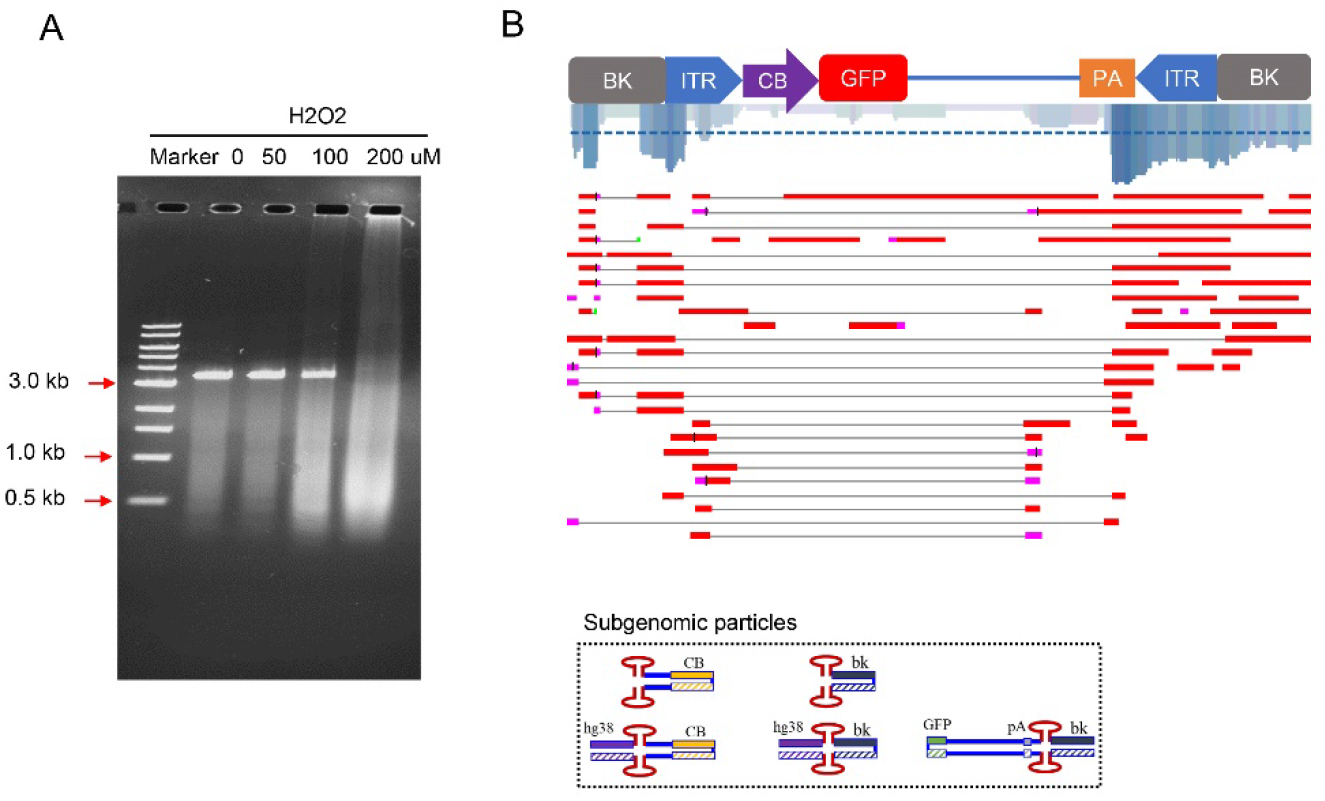
DNA damaging conditions in the host cells promoted subgenomic particle formation. Hydrogen peroxide at varying concentrations was added to the rAAV production system after transfection. A) The resulting rAAV vectors were purified by CsCl gradient, and the vector DNA analyzed by gel analysis. B) Partially recovered vector genomes were sequenced and aligned to the reference sequence. The coverage is marked by blue lines. Exemplary DNA configurations of AAV subgenomic particles are illustrated at the bottom.

## Discussion

The heterogeneity in wild type AAV virus and recombinant AAV vectors has been well documented (*3*, *9*). Similar to what has been observed for wtAAV, the subgenomic particles in rAAV vectors have similar molecular conformations: snapback genomes (SBG), genome deletion mutants (GDM), and incomplete genomes (ICG) (Fig. S1). In addition to these three major categories, a fourth category of subgenomic particles was identified as secondary derivative genome (SDG) particles arising from damage to the SBG, GDM, and IDG forms, followed by a second round of NHEJ events. The unique molecular configuration of SDG molecules prompted us to explore NHEJ events as the main cause of subgenomic DNA particle formation. While a DNA polymerase template switch mechanism may explain the formation of sSBG (*8*–*10*), the existence of GDM molecules, especially the large GDM which exceed the size of the parent AAV vector and have partial duplication of vector sequences in the junction (Fig. 1), strongly favors NHEJ as the primary mechanism that leads to the formation of subgenomic particles.

The essence of the NHEJ mechanism is the ligation of various DNA fragments. Interestingly, we were able to regenerate those SBG and GDM molecules using DNA fragments derived from AAV vector genomes, either by straight in vitro restriction endonucleases digestion or CRISPR-cas9 in vivo digestion. The generation of SBG molecules was quite efficient. Often it was the dominant vector molecules produced (Fig. 1 and Fig. 2) and supplementary Fig. S2. In addition, all features of remnant NHEJ reaction in gene editing, such as in-dels were presented in those molecules (data not shown) (*11*, *12*). Furthermore, when two fragments were introduced into the AAV packaging system, the formation of GDM could be confirmed as well (Fig. 1D and Fig. 1E). This mechanism can also explain why AAV vectors often include host genomic DNA sequence as well materials used for AAV production.

Another key point explored in this study was how the AAV fragments originated. The nickase experiment (Fig. 2) showed that simple nicking of an AAV genome was sufficient to generate corresponding SBG. Even more interesting was the identification of a stand preference for nicking. The resulting SBG contained DNA from the nicking site to its 3’ ITR. This evidence indicated that the creation of such fragments was closely coupled to DNA replication.

The nicking/lesion of DNA in the rAAV genomes suggest that any host/viral factors that were associated with AAV genomes could lead to subgenomic AAV particles formation. Hydrogen peroxide is an oxidizer that can cause global DNA damage in vivo. Our study showed that when H2O2 was present at a high concentration, rAAV production was completely ruined (Fig. 3) and resulted in the generation of primarily subgenomic particles. SBG and GDM can be observed in the sequencing analysis (Fig. S1).

Unlike the DNA template switch model (*10*), which only explains the formation of largely symmetric SBG, the NHEJ mechanism seamlessly explained the formation of SBG and GDM simultaneously. Therefore, we proposed a comprehensive model of subgenomic AAV particle formation in both wild type AAV and recombinant AAV (Fig. 4). When fragments with only one ITR are produced, it will undergo self-ligation or ligate to another fragment with only one ITR. In turn, this will create recombinant molecules with two ITRs. In case of molecules that are larger than the standard AAV size, it will not be packaged. The nicking of the AAV genome (DNA lesion) and breakage of AAV DNA leads to the formation of various DNA fragments. Both host factors and AAV proteins may cause nicking and breakage in the rAAV genomes. Ligation of these fragments generates both SBG and GMD molecules. Although it is also possible that such ligation will pick up any genomes from the host cells, the majority of the products will be SBG and GMD due to their abundance in the replication center and proximity of these fragments. The subsequent DNA replication will favor SBG or GMD when they have small genomes. Alternatively, the replication dimer of rAAV has breakage points flanking the double-D ITR, and therefore, it will efficiently self-ligate and generate SBG.

**Fig. 4.**
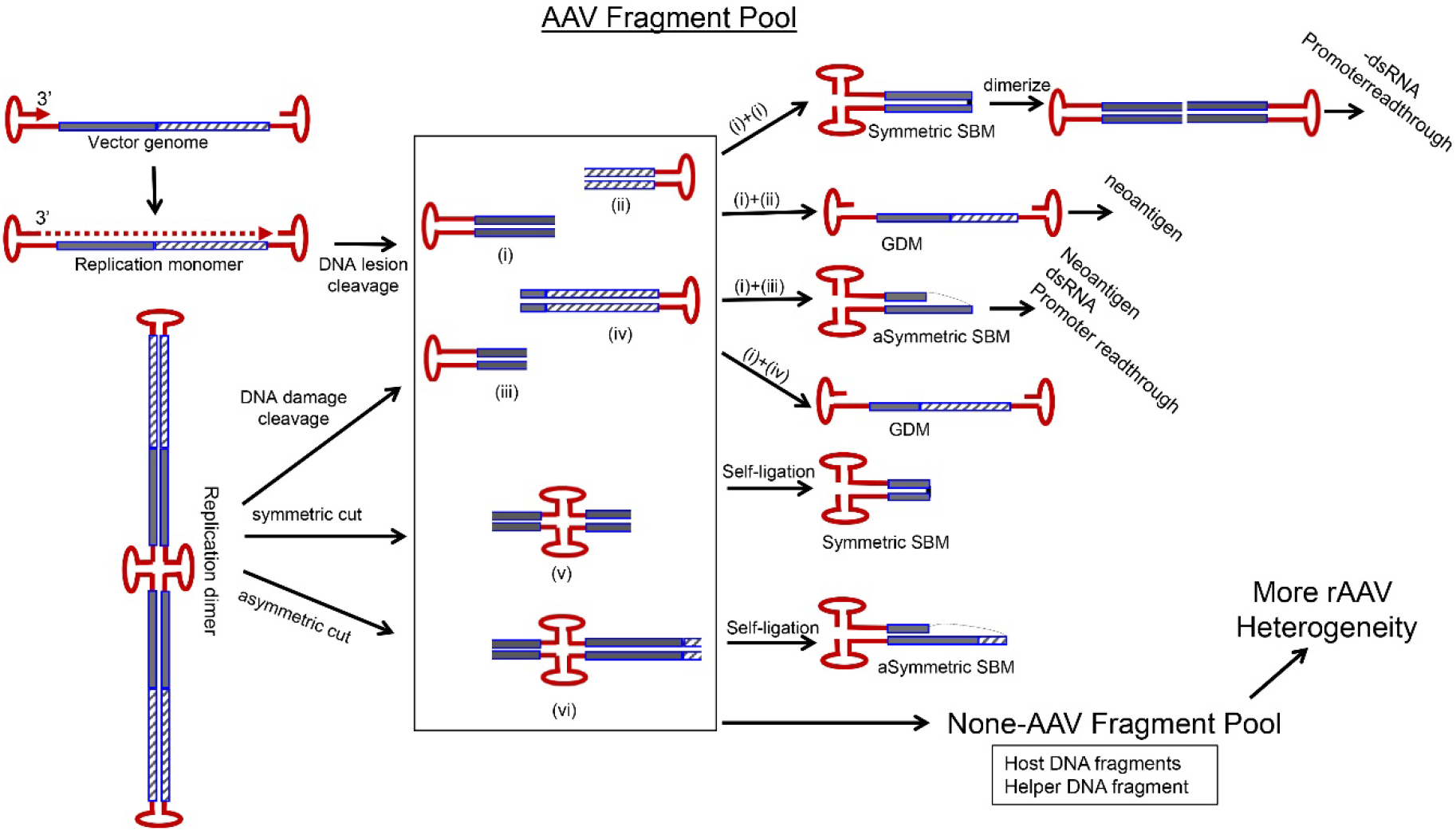
A model of subgenomic particle formation. The key point is that varying DNA fragments with only one ITR were generated from the lesion/breakage on monomer or dimer of replication form of AAV genomes. NHEJ then rejoin these fragments and the resulting products restore two ITRs in a molecule, which can be replicated and packaged in an AAV capsid. This mechanism readily led to the generation of SBG, GDM, and various forms that were not illustrated in the figure.

For wtAAV virus, subgenomic particles are beneficial for AAV life cycle (preprint bioRxiv). However, the consequences of subgenomic particles in rAAV are generally harmful. As shown in Fig. 4, subgenome particles with a promoter can produce dsRNA which will be detrimental to long term gene expression. Although dsRNA was investigated in AAV vectors (*13*, *14*), here we identified SBG as the true source for such dsRNA formation. In addition, the SBG containing only the promoter can potentially cause tumorigenesis events in the host cells or in human patients (*15*). From this study, it is clear that we need control the subgenomic particle formation in rAAV production. First, rAAV genomes should be optimized such that those sequences that are prone to breakage, i.e. nicking site for enzyme or with strong secondary structure, should be avoided. Second, host cells should be maintained in healthy condition to avoid DNA damage. Third, cellular factors should be controlled to reduce DNA damage, i.e. suppressing the factors that are part of NHEJ pathway such as ligase IV etc. Further studies should be carried out to minimize the production of SBG in rAAV vectors and improve its safety profile.

## Acknowledgments

This work was supported by grants from the National Institutes of Health (NIH) of United States (HL142019, HL114152 and HL130871).

## Supplementary Materials

Materials and Methods

Fig S1 – S2

Figure legend

