## Supplementary material for "Subgenomic satellite particle generation in recombinant AAV vectors results from DNA lesion/breakage and non-homologous end joining": Materials and Methods and Supplementary Figures

**This PDF file includes:**

Materials and Methods

Supplementary Text

Figs. S1 to S2

Materials and Methods

**Cell lines and transfection.**

HEK293 cells and GM16095 cells (a human fibroblast cell line purchased from the Coriell Institute, Camden, NJ) were cultured in DMEM supplemented with 10% fetal bovine serum, 100 μg/mL penicillin, and 100 units/mL streptomycin (Invitrogen, Carlsbad, CA). All cells were maintained in a humidified 37°C incubator with 5% CO2. PolyJet™ DNA In Vitro Transfection Reagent (SignaGen Laboratories) was used to deliver DNA into HEK 293 cells. Cells were seeded into six-well plates or 10-cm-diameter culture dishes 18 to 24 hours prior to transfection so that the monolayer cell density reached the optimal 70~80% confluency at the time of transfection. Complete culture medium with serum was freshly added to each plate 30 minutes before transfection. Prepare PolyJet™-DNA Complex for transfection according to the ratio of 3µL PolyJet™ to 1µg DNA using serum-free DMEM to dilute DNA and PolyJet™ Reagent. This was incubated for 10~15 minutes at room temperature and then the PolyJet™/ DNA mixture was added onto the medium. The PolyJet™/DNA complex-containing medium was then removed and replaced with fresh serum-free DMEM 12~18 hours post transfection.

**rAAV Infection.**

GM16095 cells were seeded into 12-well plates 24 hours prior to infection so that the monolayer cell density reached the optimal 70~80% confluency at the time of infection. The cells were washed with DMEM culture medium without serum twice, 3 min each time, before infection. 10μL of cell culture medium containing rAAV virions were added into the plate, and incubated at the indicated timepoints. GFP or mCherry fluorescence expression was observed using fluorescent microscopy (Leica D3000 B).

**Plasmid and plasmid fragments.**

Plasmid pH22 was the helper plasmid used that contained the rep and cap coding sequences. Plasmid pFd6 was the miniadenovirus helper. Plasmid pCB-GFP-3.4K contained a cytomegalovirus enhancer and beta -active promoter. This plasmid was used to make vector containing the green fluorescent protein (GFP) reporter gene flanked by the AAV ITR.

Plasmid pCB-GFP-6.4K was made by cloning a 3kb stuff DNA into pCB-GFP-3.4K. The pCB-GFP-6.4K plasmid was subjected to a series of restriction digestion to produce a series of DNA fragments with different lengths, that where, fCB-GFP-0.6K, fCB-GFP-1.0K, fCB-GFP-1.6K, fCB-GFP-1.8K, fCB-GFP-2.3K and fCB-GFP-3.1K.

The gRNA target sequences in the pCB-GFP-3.4K rAAV genome were designed using the Broad Institute gRNA designer tool (https://www.broadinstitute.org/rnai/public/analysis-tools/sgrna-design). The sequences for these sgRNA targets are gRNA 4,GGG AGC GGG ATC AGC CAC CG, gRNA 5:AAG CTG CGG AAT TGT ACC CG, gRNA 9:TTA GTC GAC CTC GAG CAG TG, gRNA 10:TGT TCC GGC TGT CAG CGC AG, gRNA 13:GAT CAG CGA GCT CTA GTC GA

**Virus production and purification.**

AAV viruses were produced using the triple plasmid transfection system in HEK 293 cells. PolyJet™ DNA In Vitro Transfection Reagent (SignaGen Laboratories) was used to deliver DNA into the HEK 293 cells. At 72 hours after transfection, medium was collected and precipitated with 40% of PEG (finial concentration 8%) overnight at 4°C. Next, it was centrifugated, resuspended and treated with DNaseI. AAV of different densities were separated using CsCl gradient ultracentrifugation. AAV of different densities were extracted and dialyzed against 5% sorbitol in Phosphate Buffered Saline (PBS, NaCl 137 mM, KCl 2.7 mM, Na2HPO4 10 mM, KH2PO4 1.8 mM, pH 7.2). Vector genome titers were determined by quantitative real-time PCR (qPCR), with vector titers expressed as vg/ml. To obtain vectors representative of all viral particles, the gradient centrifugation step was skipped. Three days after transfection, the medium was collected and precipitated into concentrated solution of rAAV particles. rAAV genomic DNA was purified and further analyzed using agarose gel electrophoresis and qPCR.

**DNA agarose gel electrophoresis.**

rAAV genome was extracted and purified as followed: Viral vectors were treated with DNase I (1U/mL) for 30 min at 37°C, then 1 µl of 0.5 M EDTA was added (to a final concentration of 5 mM) and subsequently heated for 10 min at 75°C to cease DNase I activity. 1/2 volume of lysis buffer (Direct PCR Tail, Viagen) containing proteinase K (40 μg/mL) was added and incubated for 1 hour at 56°C and finally heated for 10 min at 95°C. One volume of phenol: chloroform: isoamyl alcohol (25:24:1) was added to the samples, and vortexed thoroughly for approximately 20 seconds. They were then centrifuged at 4°C for 30 minutes at 16,000 × g. The upper aqueous phase was carefully removed and transferred to a fresh tube. 200 μL of 70% ethanol was added, and the tubes centrifuged at 4°C for 10 minutes at 16,000 × g. The supernatant was carefully removed and the pellet was allowed to air dry at room temperature. 20ul of TE buffer was added to dissolve DNA. DNA concentration was measured using Nanodrop. 100ng DNA was loaded on 1% of Neutralizing gel and run at 120V for 50min. The equal DNA was loaded on a 1% of Alkaline gel and run at 60V for 100min in ice-water bath. Gels were stained using 1×SYBR@ Safe DNA gel stain (Invitrogen) and a photo taken at a wavelength of 365nm using a ChemiDOCTMMP Imaging System (Bio-rad).

**H2O2 Treatment.**

HEK 293 cells were seeded in twenty 15-cm dishes and incubated for 18 hours. The old culture medium was replaced with free FBS DMEM containing a final concentration of 0µM, 50µM, 100µM and 200µM H2O2 at 60 min prior to transfection. Three plasmids, pH22, pFΔ6 and pssAAV-CB-GFP-4.7K, were transfected into HEK 293 cells using PolyJet™ DNA In Vitro Transfection Reagent (SignaGen Laboratories). 72 hours after transfection, medium was collected, precipitated with 40% of PEG (finial concentration 8%), and purified by Cscl gradient method. rAAV DNA was extracted and purified using the phenol: chloroform: isoamyl alcohol (25:24:1) method. 500ng of rAAV DNA was subjected to sequence by PacBio SMRT platform, meanwhile, 30ng of DNA was loaded on 1% agarose gel and run at 120V for 50min.

**Quantitative Real Time PCR (qPCR) Assay.**

Viral vectors (1 × 1010 vg, 1ul) in solution containing DNase I (1U/mL) were incubated for 30 min at 37°C, add 1 µl of 0.5 M EDTA (to a final concentration of 5 mM) and subsequently heated for 10 min at 75°C to cease DNase I activity. Control samples each received lysis buffer (Direct PCR Tail, Viagen) containing proteinase K (40 μg/mL), and were incubated for 1 hour at 56°C and finally heated for 10 min at 95°C. The samples intended for thermal treatment were directly heated following heat inactivation of DNase I treatment at the indicated temperatures. The copy numbers of viral genomes subsequently released were quantified by real-time PCR and expressed in vg/ml. The primers include GFP forward:AGTCCGCCCTGAGCAAAGAC and GFP reverse:CTCGTCCATGCCGAGAGTGA; polyA forward:GTGCCTTCCTTGACCCTGGA and polyA reverse: CACCTACTCAGACAATGCGATGC.

**AAV Genome Sequencing and Data Analysis.**

For long-read PacBio SMRT sequencing, AAV samples were prepared according to SMRTbellTM procedures. DNA was extracted and purified by AMPure PB Beads and then repaired by SMRTbellTM Damage Repair kit. The adaptor ligation reaction was performed, and then ExoIII and ExoVII were added to remove failed ligation products. AMPure PB were performed three times.

SMRT subread filtering and the high-quality circular consensus sequences corresponding to the rAAV library were generated using SMRT analysis portal (minimum accuracy of 0.99 and minimum of 3 CCS passes), and considered for further analysis. Filtered reads were mapped to the rAAV genome using the Minimap2 (https://github.com/PacificBiosciences/pbmm2) and processed alignments to demonstrate configuration categories of molecules in the rAAV population.

Supplementary Text

Fig. S1.

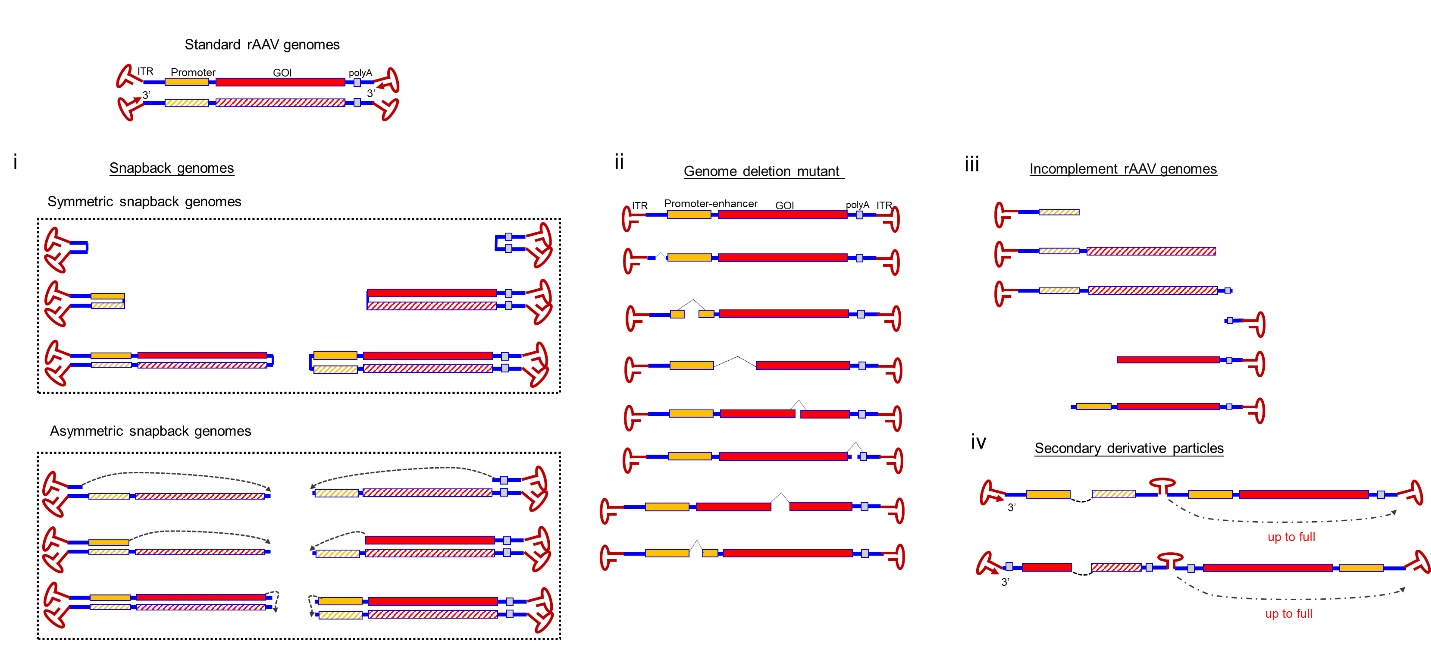

Fig. S1. Molecular configuration of DNA genomes in rAAV vectors. AAV genomes were sequenced using the PacBio SMRT platform and compared to the reference sequences on the top. Besides the standard sized AAV vector genomes, four typical categories of subgenomic rAAV genomes were found in the rAAV vectors: i). symmetric snapback genomes (sSBG) and asymmetric snapback genomes (aSBG); ii). Genome deletion mutants (GDM); iii). Incomplete genomes (ICG); iv). Secondary derivative genomes (SDG).

Fig. S2.

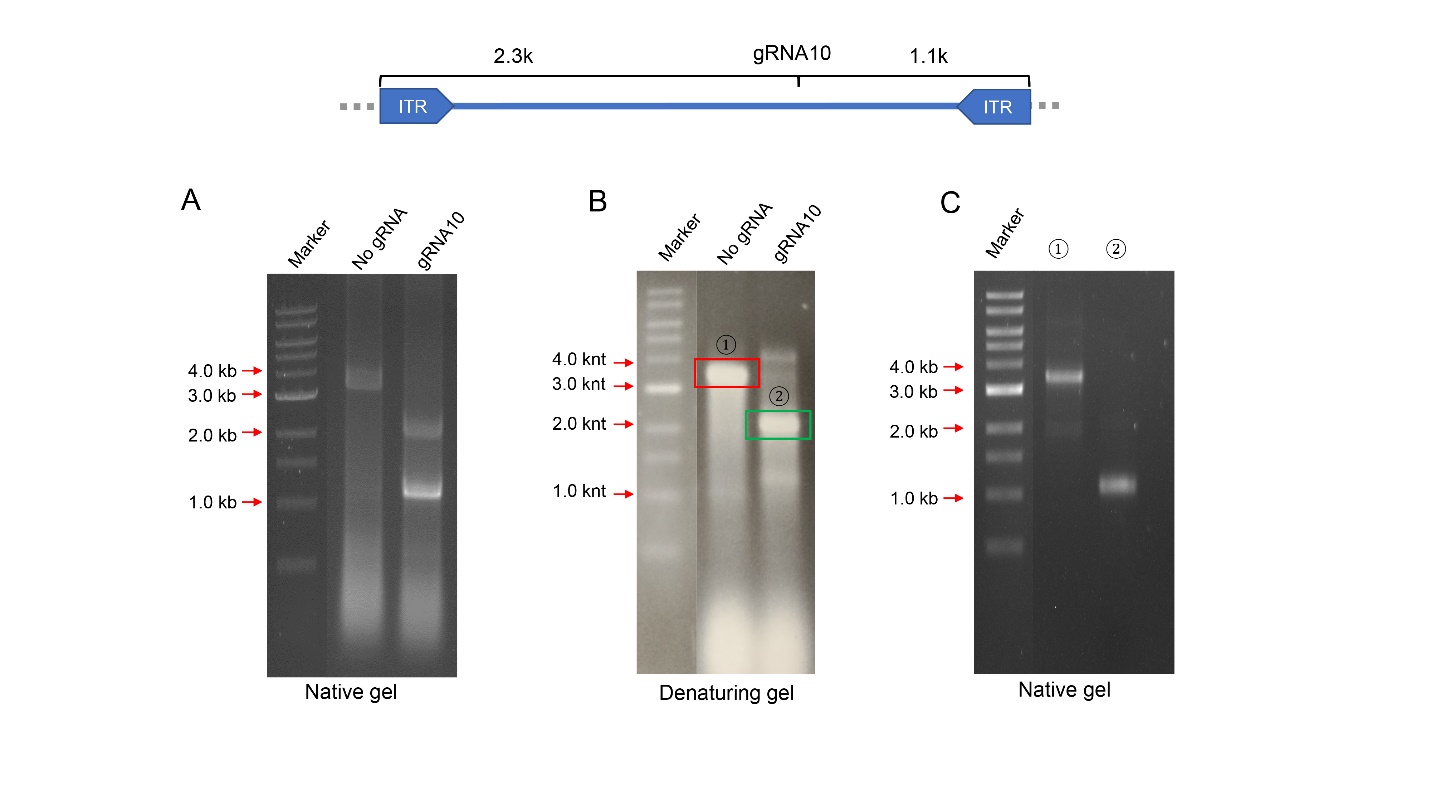

Fig. S2. Intra-host cell vector DNA breakage is the mechanism for AAV subgenomic particle formation. Hek293 cells were transfected with AAV plasmid pCB-GFP-3.4k along with Cas9 expressing plasmids with or without corresponding guide RNA. A) The resulting vector DNA was electrophoresed in native agarose gel. B) The resulting vector DNA was electrophoresed in denaturing agarose gel. C) The denatured fragments of B (indicated as ① ②) were collected and renatured and were electrophoresed again in the native gel.

We then created an in vivo model using the CRISPR-Cas9 system to mimic the breakage that occurred in vivo. In the vector production system, the Cas9 expression plasmid was included with the vector plasmid pCB-EGFP-3.4k. In contrast to the control without guide RNA, transfection with guide RNA produced two distinct vectors with a size of 2.3kb and 1.1kb in a native gel (Fig. S2A), which corresponded to the cutting site. In the denaturing gel, the original vector was present as a 3.4kNT single-stranded DNA. In contrast, the vectors produced in the presence of guide RNA appeared as single stranded DNA that were 4.6kb or 2.2kb in size. We then recovered the 2.2 knt single stranded DNA from the gel, renatured the DNA, and electrophoresed it in the native gel. The 2.2kNT DNA fragment appeared as 1.1kb dsDNA, which were confirmed by restriction digestion (data not shown). These results suggested that the SBG molecules for either direction were formed in the presence of CRISPR-Cas9 induced digestion in vivo (Fig. S2C).
